# Polygenic and redundant architectures of climate-adaptive traits may complicate genomic predictions of maladaptation

**DOI:** 10.64898/2026.02.02.703187

**Authors:** Rafael Candido-Ribeiro, Brandon M. Lind, Pooja Singh, Mengmeng Lu, Dragana O. Vidakovic, Christine Chourmouzis, Tom R. Booker, Sam Yeaman, Sally N. Aitken

**Affiliations:** Department of Forest and Conservation Sciences, University of British Columbia, Vancouver, BC, Canada V6T 2G9; Department of Ecology & Evolutionary Biology, and Institute for Systems Genomics, University of Connecticut, Storrs, CT 06269, USA; Department of Biological Sciences, University of Calgary, Calgary, Canada T2N 1N4; Aquatic Ecology & Evolution Division, Institute of Ecology and Evolution, University of Bern, CH-3012, Switzerland; Department of Fish Ecology & Evolution, Swiss Federal Institute of Aquatic Science and Technology (EAWAG), Kastanienbaum, CH-6047, Switzerland; Department of Biological Sciences, University of Notre Dame, Notre Dame, IN 46556, USA

**Author notes:** Corresponding authors: Rafael Candido-Ribeiro and Sally N. Aitken, **Email:** &.

**Keywords:** genomic prediction, polygenic traits, local adaptation, case-control genome-wide associations, genetic redundancy

## Abstract

**Significance:** Understanding the genetic basis of local adaptation is an important step to predict species’ responses to climate change. Current predictions of maladaptation to climate in trees, however, rely mostly on genotype-environment associations, which overlooks the complexity of genetic architectures underpinning climate-adaptive traits. Based on genotype-phenotype associations, we unveil key aspects of the polygenic architecture of drought tolerance and cold hardiness in two lineages of a widespread conifer. Our results suggest that predictions based solely on climate-associated alleles may mischaracterize maladaptation to future climates due to mismatches between the distribution of climate-associated alleles and alleles contributing to adaptive traits. We also show that cold hardiness and drought tolerance evolved largely independently in the two lineages, and with little pleiotropy between traits.

Understanding the genetic architecture of climate-adaptive traits can help predict responses to climate change and inform management strategies that mitigate the effects of future climates. By characterizing the genetic basis of adaptation to spatially varying climate conditions, models can predict response to temporal change in climate by extrapolating from spatial patterns of variation. However, such methods are predicated upon a simple genetic basis, with a linear relationship between allele frequency, phenotype, and environment. Here, we explore the nature of the mapping between genotype, phenotype, and climate adaptation in Douglas-fir, an important North American conifer. Using two large common garden experiments combined with exome sequencing and case-control genome-wide association tests, we found that drought tolerance and cold hardiness in two Douglas-fir varieties (*Pseudotsuga menziesii* var. *menziesii* and var. *glauca*) are highly polygenic with little pleiotropy between the two traits. The two varieties showed little repeated evolution in the genetic basis of drought tolerance and none for cold hardiness. Allelic clines were observed along geographic and climatic gradients across 74 natural populations, but their direction of effect was often inconsistent with phenotypic clines. Our findings reveal that the genomic variation underlying climate-adaptive traits is remarkably complex, with redundancy likely playing a key role in the evolution of the divergent polygenetic architectures of abiotic stress tolerance. This suggests that predicting maladaptation to climate based solely on climate-associated alleles may misrepresent future adaptive responses. Genomic predictions and management practices should focus on individual lineages and consider the particularities of the genetic architecture of climate-adaptive traits.

## Introduction

Growing climate variability caused by anthropogenic impacts poses serious risks to ecological communities and the ecosystem services they provide (1, 2), and is expected to increase the maladaptation of natural populations of trees as climates deviate from the historical conditions that shaped current patterns of adaptation (3). In temperate forests, cold temperatures have historically driven much of the local adaptation in trees, in particular conifers (4, 5). With warming, other climatic selection pressures will likely rise in importance. For example, extreme droughts are becoming more frequent and severe (6). On the other hand, with increasing variability in climate, frost events are also expected to become more unpredictable, and adaptations to freezing temperatures will remain crucial (7–10).

For plants evolving in response to a fast climate change scenario, drivers of selection are expected to act primarily upon standing genetic variation present across populations (11, 12), in particular on variation that is already adaptive (13). Studying how complex climate-adaptive traits may be affected by the number of underlying loci, their effects sizes and allele frequencies, as well as the levels of genetic redundancy and pleiotropy (i.e., genetic architecture, 14) present in different populations is key to help predict the adaptive potential of trees to future climatic conditions (15), inform management strategies that mitigate maladaptation to future climates (e.g., breeding, assisted gene flow, 16), and ultimately, further elucidate evolution in long-lived organisms (17). An increasingly popular approach in landscape genomics is to model the distribution of genotype-environment-associated alleles (GEA; reviewed in ref. 18) across environmental variation in both space and time to predict future maladaptation (e.g., ref. 19 and 20). These predictions assume the identified loci affect traits relevant for adaptation to future climates (21). The capacity for detecting adaptive variants associated with climate can vary depending on the genetic architectures controlling the traits involved in local adaptation (22). However, GEA approaches do not use phenotypic data from common garden experiments (23), which makes it difficult to validate the relationship between the identified variants and the traits contributing to local adaptation (15, 22). Additionally, GEA methods can generate large numbers of false positives when populations are distributed across environmental gradients that covary with the demographic history of the species (24), or of false negatives when models are over-corrected for the same issue (25). Identifying and using loci directly associated with climate-adaptive traits to test hypotheses of ecological importance (e.g., local adaptation across environments) and in spatiotemporal predictions of maladaptation is expected to produce more robust results than GEA (22, 26).

Key questions for understanding the evolutionary mechanisms underlying climate adaptation in trees include whether complex climate-adaptive traits are affected by the same alleles globally (i.e., across populations), and whether different alleles and genetic combinations are able to produce similar phenotypes and physiological responses locally in different populations (i.e., genetic redundancy; 27). Genetic redundancy is expected for polygenic adaptive traits (28), and is predicted to constrain allelic clines for traits associated with the environment (29), further complicating the use of genotype-environment associated loci in studies of local adaptation. However, this still needs to be empirically explored. Therefore, clines in allele frequencies along climatic gradients expected to drive selection should be tested for loci underpinning climate-adaptive traits (26, 29).

A certain degree of repeated evolution (i.e., overlap in genes affecting traits involved in adaptation) has been observed between related taxa exposed to similar selection pressures (30–32), such as climate (e.g., 33). Phylogenetically closer taxa are expected to show higher levels of repeated adaptation (32, 34), but highly polygenic traits are less likely to show repeatability in adaptive genetic responses to climate (17). On the other hand, pleiotropy can play a role in climate adaptation in conifers (35), and may facilitate repeated evolution (36). Pleiotropic genes controlling for responses to drought tolerance and cold hardiness, for example, have been observed in some groups of plants (37, 38); however, the extent to which pleiotropic genes interact and affect climate-associated traits in conifers is still largely unknown.

By looking at the distribution of putatively adaptive genetic variation, it is possible to infer which populations may be better prepared for future climates and extreme events, and to identify potentially useful genetic variation for breeding or translocations. Genetic variation for drought tolerance in seedlings of the two Douglas-fir varieties (*Pseudotsuga menziesii* var. *menziesii*, and var. *glauca*) is primarely maintained within populations, with weak to absent signals of local adaptation to drought within varieties (39), and a narrow-sense heritability (h^2^) of *c*. 0.2 (40). A much stronger signal of range-wide local adaptation to freezing conditions was observed within varieties (41, 42), but a considerable proportion of this variation is also maintained within populations, with larger h^2^ (*c.* 0.4, ref. 40). The two Douglas-fir varieties, which diverged c. 2.1 million years ago (43), experience similar challenges such as droughts and freezing events to varying degrees, despite of the lack of overlap in their geographic ranges. As a result, some repeatability in the genetic basis of drought tolerance and cold hardiness is expected, but little information exists on the extent of adaptive repeatability between closely related conifer taxa.

Here, we explore the within-population genomic variation associated with tolerance to drought and freezing temperatures using case-control genome wide association (GWAS) approaches in 20 natural populations from across the geographic ranges of the two main varieties of Douglas-fir (Fig. 1), spanning a large gradient of climatic conditions (*SI Appendix,* Fig. S1), to identify candidate genes important for climate adaptation. We then tested for the overlap of candidate genes between traits and between varieties and investigated clines in the frequencies of trait-associated alleles along climate gradients of 74 independently genotyped natural populations. Specifically, we ask: 1) how similar are the genetic architectures underpinning drought tolerance and cold hardiness between the two Douglas-fir varieties? 2) What is the extent of pleiotropy for drought tolerance and cold hardiness within varieties? 3) Are the same climatic drivers of local adaptation identified using genomic and phenotypic data? Finally, 4) If allelic clines are present, are they localized or distributed range wide, and will they be on par with phenotypic clines?

**Fig. 1.**
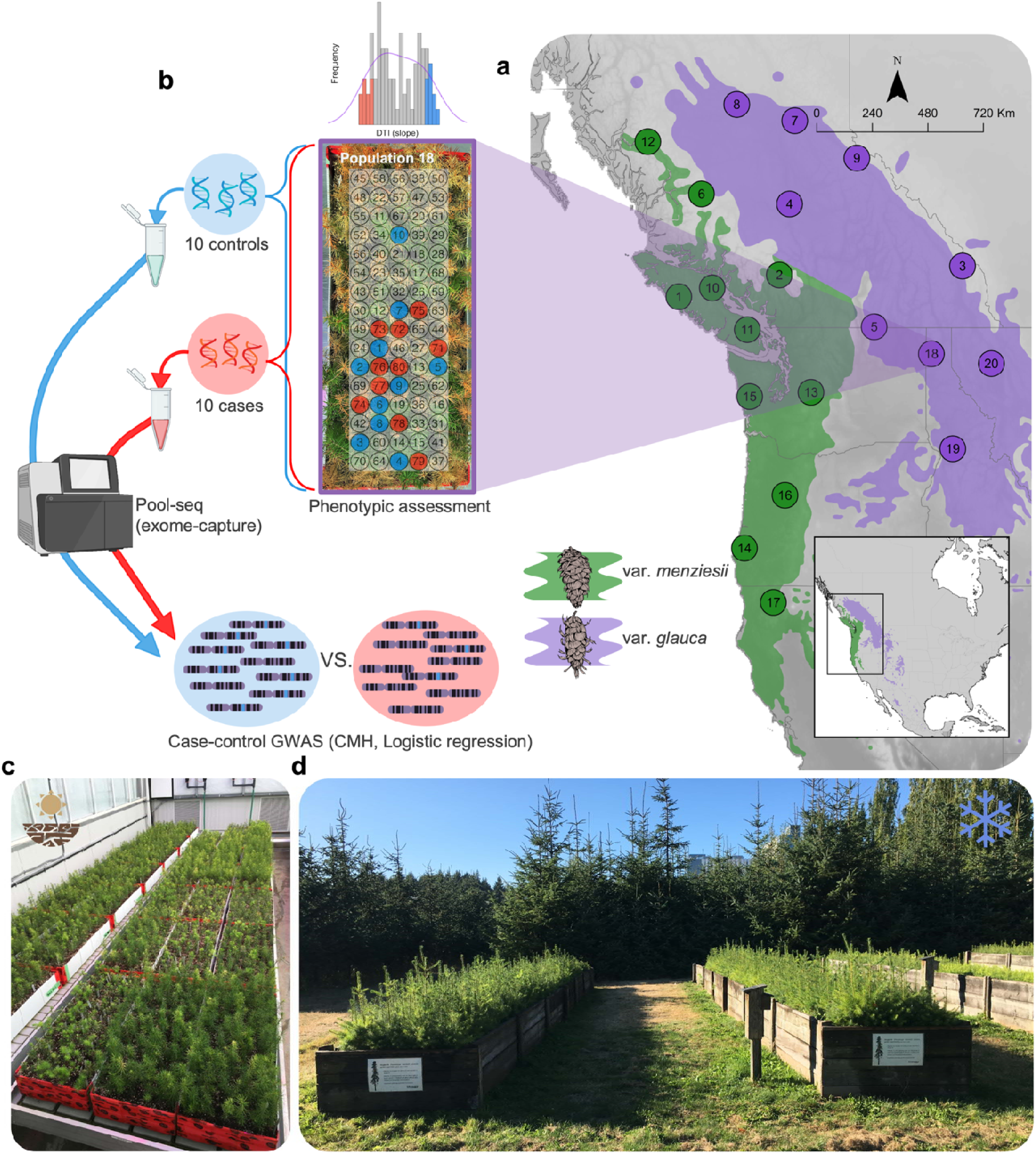
Distribution of the 20 sampled natural populations of Douglas-fir for the case-control GWAS and example of workflow for the drought tolerance study for population 18 **a**) Range map with the selected populations of var. *glauca* and var. *menziesii*. **b**) Example of workflow for the study: ranking individuals for drought tolerance (DTI) (distribution on top) after the drought-to-death treatment; selection of cases (least tolerant, in red) and controls (most tolerant, in blue) within one population (population 18, var. *glauca*); DNA extraction and pooling, pool-sequencing; and association test approaches (GWAS)–CMH (Cochran-Mantel-Haenszel test) and logistic regression (with a quasi-binomial distribution). The sampling strategy for cases and controls in the cold hardiness experiment was similar (see Methods). All 20 case-control populations were tested for drought tolerance (**c**) and cold hardiness (**d**) in separate experiments. The numbers 1-80 inside the box in **b** indicate the ranking of individuals after phenotyping for DTI from most to least tolerant in population 18.

## Results

### Complex genetic architectures underpinning drought tolerance and cold hardiness in Douglas-fir

By assessing the phenotypic variation in seedling drought tolerance and cold hardiness within 20 natural populations of Douglas-fir (*Pseudotsuga menziesii*) (*SI Appendix*), planted in two large common garden experiments (3,433 seedlings), we selected the ten least (cases) and the ten most tolerant individuals (controls) for each trait within each population (i.e., extreme phenotypes; *SI Appendix*, Fig. S2, S3). We observed substantial phenotypic variation within populations for both traits, with significant differences between cases and controls (*p-value* < 0.001), partially confirming the results of larger variation within than among populations, observed by 39.

Next, we established DNA pools of cases and controls with approximately equal DNA contribution per individual and used targeted exome capture of each population and pooled-sequencing (*SI Appendix*). A different number of SNPs was obtained for each dataset (i.e., variety x trait combination) after SNP calling and filtering (44; *SI Appendix*). All retained SNPs were located within the transcribed region of a gene (hereafter called gene) (*SI Appendix*, Table S2). The var. *menziesii* datasets consisted of 439,047 and 196,946 SNPs for the drought tolerance and cold hardiness traits, respectively. The var. *glauca* datasets consisted of 637,491 and 708,369 SNPs for drought tolerance and cold hardiness, respectively.

We then considered the overlap between two case-control GWAS approaches–CMH (45; *q-value* ≤ 0.05) and logistic regression (46) *p-value* ≤ 0.05)–per dataset to determine the SNPs significantly associated with each trait. This was done to take advantage of the power of CMH tests to detect associated SNPs, but also reduce the number of false positives produced by this method by validating with the logistic regression approach (see *SI Appendix* for details).

The large within-population variation for drought tolerance and cold hardiness in both Douglas-fir varieties, combined with the moderate heritability observed for both traits in this species (40), enabled the detection of hundreds of loci highly associated with both traits. We obtained 303 SNPs and 51 SNPs for var. *menziesii* drought tolerance and cold hardiness, respectively; and 393 SNPs and 371 SNPs for var. *glauca* drought tolerance and cold hardiness, respectively (Fig. 2; See *SI Appendix*, Table S2 for the number of SNPs with significant but non-corrected *p-values*). Highly polygenic architectures are common for complex adaptive traits in plants (47–50), but the genetic basis of adaptation in complex traits may also be oligogenic (51) or may involve selective sweeps in one or few loci (52).

**Fig. 2.**
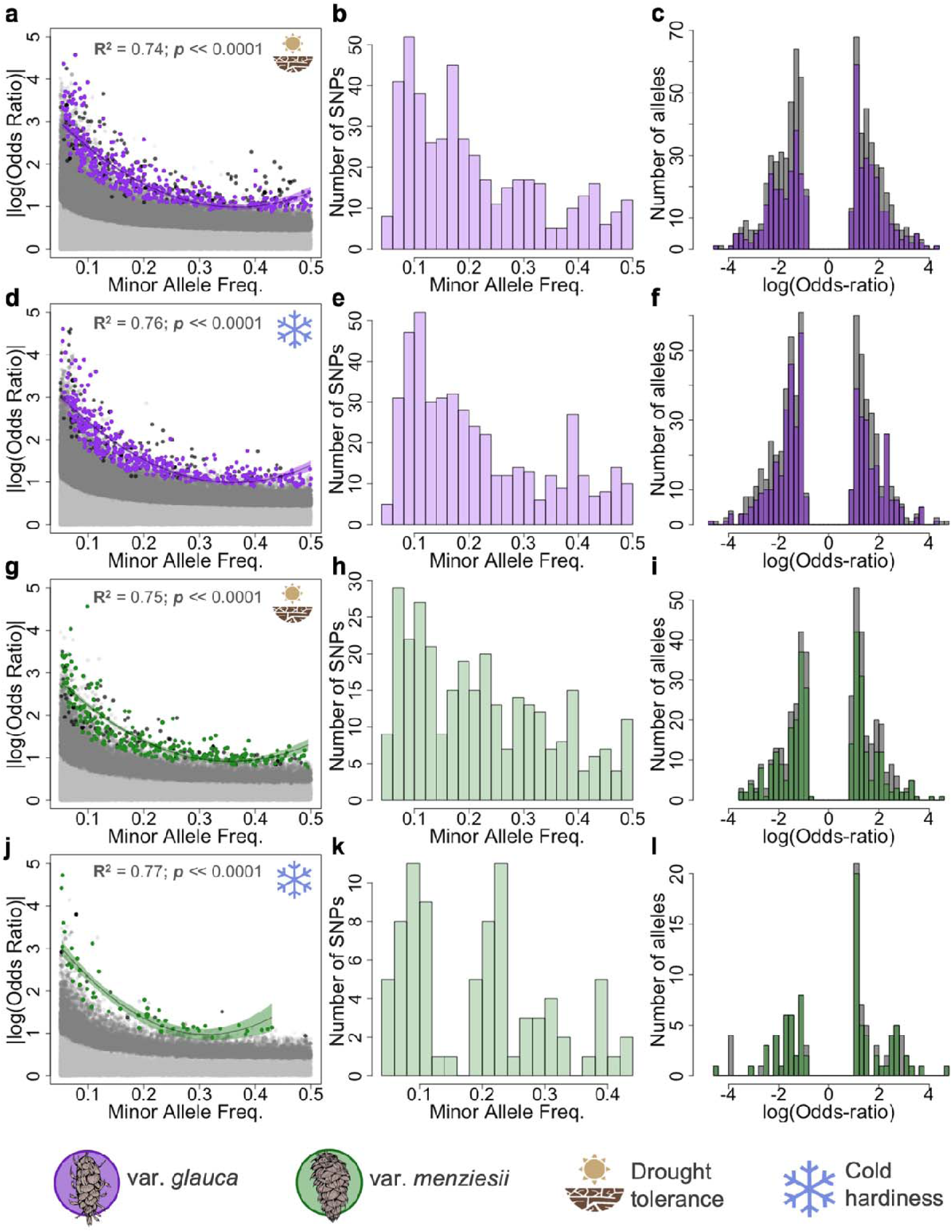
Significant SNPs associated with drought tolerance and cold hardiness for var. *glauca* and var. *menziesii*. Relationship between effect size (|log(Odds-ratio)|) and minor allele frequency (MAF) for SNPs significantly associated with the traits drought tolerance (**a** and **g**) and cold hardiness (**d** and **j)**. Colored points (purple for var. *glauca* and green for var. *menziesii*) are based on the overlap of Cochran-Mantel-Haenszel (CMH) test significant SNPs corrected for false discovery rate (*q-value* ≤ 0.05) and the quasi-binomial logistic regression significant SNPs (*p-value* ≤ 0.05). Light-gray points represent ∼20,000 random non-significant SNPs. Dark-gray points represent CMH significant SNPs before correction for FDR (*p-value* ≤ 0.05). Black points represent CMH significant SNPs after FDR correction (*q-value* ≤ 0.05). **b**, **e**, **h**, and **k**) distribution of minor allele frequencies for the associated SNPs. **c**, **f**, **i**, and **l**) distribution of log(Odds-ratios) from the CMH tests solely corrected for FDR (gray), and that overlap with the significant quasi-binomial logistic regression SNPs (colored). MAFs used here are among the 20 case-control populations.

To verify the potential polygenic nature of drought tolerance and cold hardiness–in which the majority of alleles controlling these traits are expected to have small effect sizes and higher frequencies–the relationship between allele frequencies and allele effect sizes was assessed. We considered the log(Odds Ratio) from the CMH tests as the approximate effect size of the allele on the tested trait. SNPs with larger effect sizes (|log(Odds Ratio)|) were observed at intermediate minor allele frequencies (MAF = 0.05-0.2) for both drought tolerance and cold hardiness in both varieties (Fig. 2a d, g, j). Similarly, a larger number of significant SNPs was detected at intermediate minor allele frequencies (MAF = 0.05-0.2) than at higher frequencies (MAF > 0.2; Fig. 2b, e, h, k).

Using the non-parametric Wilcoxon Sum test, we tested whether the differences in average allelic effect sizes (|log(Odds ratio)|) observed between varieties and between traits within varieties were significant. Within var. *menziesii*, a lower number of loci, but with larger effect (|log(Odds ratio)| = 1.83) contributed to cold hardiness on average than to drought tolerance (|log(Odds ratio)| = 1.52; *p-value* = 0.005). Within var. *glauca*, no significant difference in effect sizes was observed between the two traits (cold hardiness (|log(Odds ratio)| = 1.75; drought tolerance (|log(Odds ratio)| = 1.78). The average effect size of var. *glauca* drought tolerance SNPs was significantly larger than var. *menziesii*’s (*p-value* << 0.001) (*SI Appendix*, Fig. S4).

### Lack of repeated evolution between varieties and a small signal of pleiotropy between traits

After mapping the significant drought tolerance SNPs (corrected for false discovery rate (FDR)) to the annotated Douglas-fir genome, 266 genes with at least one significant SNP per gene (*q-value* ≤ 0.05) were obtained for var. *menziesii* and 317 genes for var. *glauca*. The cold hardiness SNPs were mapped to 53 genes for var. *menziesii*, and 314 genes for var. *glauca* (Fig. 3b). All candidate genes were tested for GO term enrichment. In var. *glauca*, candidate genes for drought tolerance were significantly enriched for GO terms related to abscisic acid metabolism including abscisic acid catabolic process and (+)-abscisic acid 8’-hydroxylase activity. In var. *menziesii*, enriched GO terms included positive regulation of stomatal complex development and stomatal complex development, both of which are known to be involved in drought responses. For candidate genes for cold hardiness, var. *glauca* showed enrichment in GO terms related to regulation of fatty acid oxidation, regulation of lipid catabolic process, and regulation of photorespiration, while var. *menziesii* exhibited enrichment in UDP-glucose metabolic process and lipid glycosylation (see *SI Appendix*, Table S14, S15, S16, and S17 for the full list of significant GO terms).

**Fig. 3.**
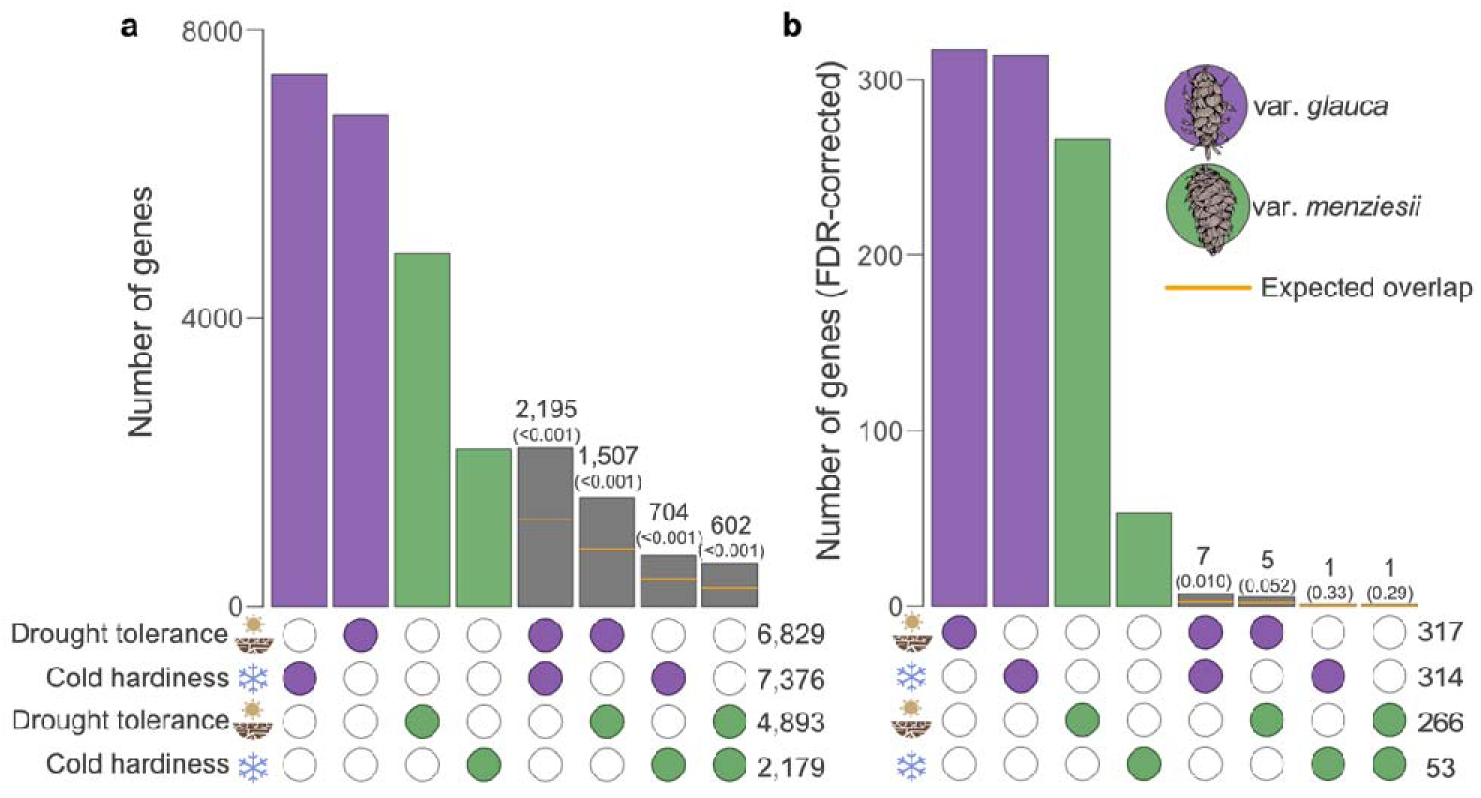
Number of drought tolerance and cold hardiness candidate genes identified for var. *glauca* and var. *menziesii* and the overlaps between traits and between varieties. Candidate genes in **a** contain at least one mapped significant SNP (*p-value* ≤ 0.05, not corrected for false discovery rate) from the intersection between the CMH and logistic regression GWAS tests for each combination of trait and variety. Candidate genes in **b** contain at least one highly significant SNP from the intersection between the CMH test (*q-value* ≤ 0.05, corrected for false discovery rate) and logistic regression test (*p-value* ≤ 0.05) for each combination of trait and variety. Gray bars depict the number of overlapped genes between groups. Numbers in parenthesis depict the *p-value* of the hypergeometric test used to determine the significance of the overlap.

Annotated candidate genes implicated in drought stress response include CCD1, and CHLH in var. *glauca*, and ALDH2C4 and CKX5 in var. *menziesii*. Genes previously known to be involved in cold hardiness were also annotated, such as CIPK20, and UBC13 in var. *glauca*, and HT1, and AAE7 in var. *menziesii* (*SI Appendix*, Tables S10, S11, S12, S13).

To infer the potential repeated adaptation to cold temperatures and drought between the two varieties, as well as pleiotropy between climate-adaptive traits within each variety, we tested the significance of the number of overlapping candidate genes between groups using a hypergeometric distribution. Seven genes overlapped between drought tolerance and cold hardiness in var. *glauca*, which was significantly higher than the 2.4 expected overlap based on the hypergeometric test (*p-value* = 0.010). No overlap of genes was observed between traits in var. *menziesii* (Fig. 3b). With a marginal significance, only five drought tolerance genes overlapped between var. *menziesii* and var. *glauca* (*p-value* = 0.052, expected overlap = 2; Fig. 3b). As an alternative approach to identify genes driving trait variation, we compared the observed number of trait-associated SNPs per gene with the binomial expectation based on the total number of SNPs per gene (*i.e*., the 0.999 quantile of the binomial distribution for each trait and variety, as described in ref. 33, and again, no significant overlap was observed in the most associated genes for the different traits. Overall, these results suggest that the two Douglas-fir varieties have adapted largely independently to drought and freezing conditions and that there is not much evidence for pleiotropy between drought tolerance and cold hardiness in Douglas-fir. For the number and overlap of genes with non-FDR-corrected SNPs, see *SI Appendix*.

### Substantial population differentiation within var. *glauca* for cold hardiness, and between varieties for drought tolerance and cold hardiness

To evaluate evidence for local adaptation at the trait and locus level, we explored patterns of differentiation among populations and linkage disequilibrium (LD) using an additional dataset, calling SNPs from an expanded pool-seq dataset encompassing 74 populations (one pool of *c*. 40 individuals per population) from the natural range of Douglas-fir (Fig. 4i; *SI Appendix*, Fig. S1; 53). At the phenotypic level, we reported the phenotypic differentiation (V) among populations and between varieties for each trait (as described in ref. 54), and qualitatively compared them with their respective background and trait-associated *F_ST_* estimates. Except for the population differentiation for drought tolerance within var. *menziesii* and species-wide, all estimates of *V_pop_* were larger than *F_ST_* for associated and background SNPs (Table 1), showing that the phenotypes measured in our experiments are exhibiting evidence consistent with local adaptation. Phenotypic differentiation for cold hardiness, for example, was *c.* 12 times larger than the background *F_ST_* in var. *menziesii*, and *c.* 11 times larger in var. *glauca*.

**Fig. 4.**
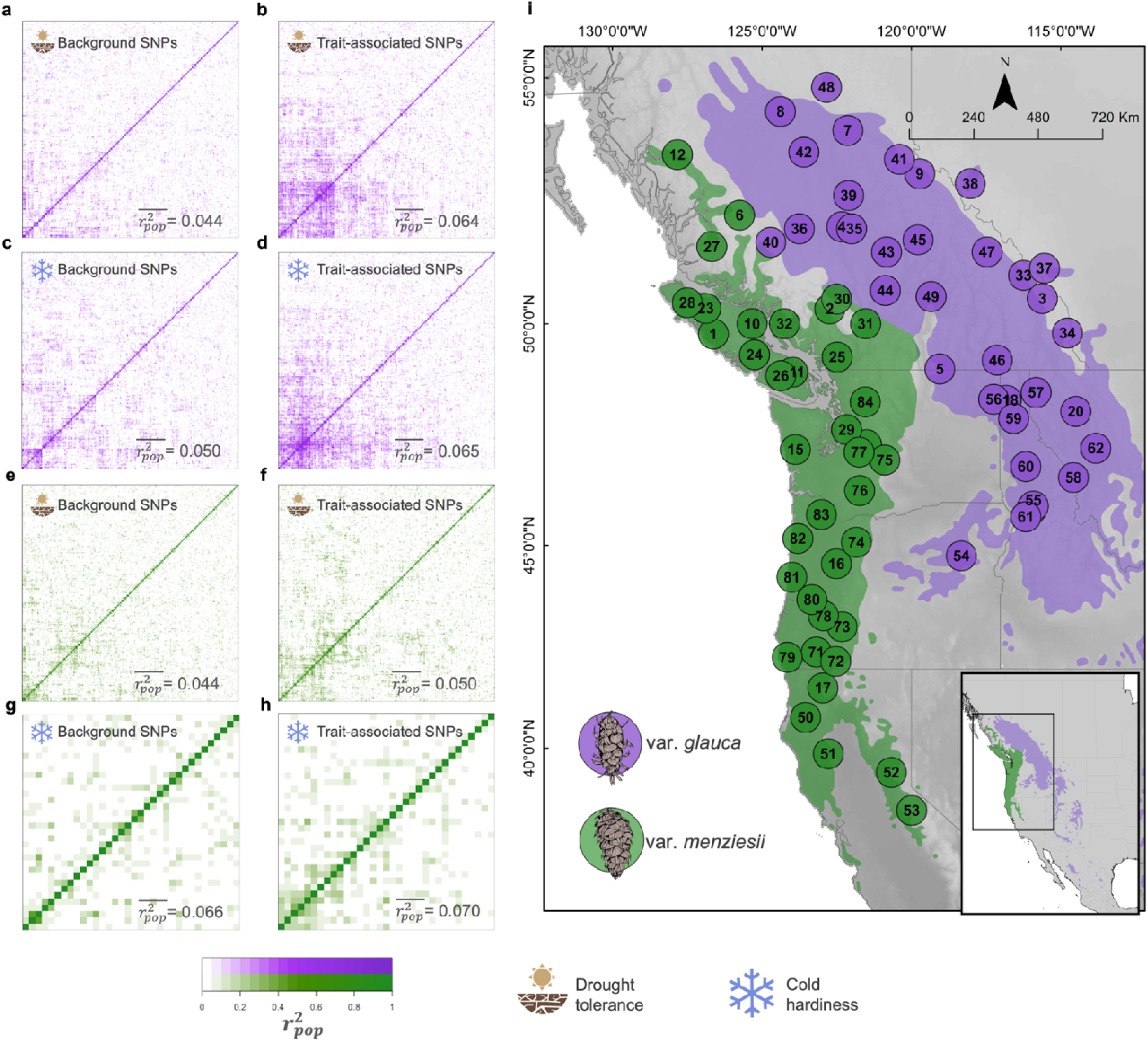
Pairwise linkage disequilibrium (LD,) for each Douglas-fir variety with SNPs called for 74 natural populations (35 var. *glauca*, and 39 var. *menziesii*) from across the two varieties’ ranges (**i**). Pairwise LD () across Douglas-fir natural populations for random background SNPs (**a**, **c**, **e**, **g**) and top significant (**b**, **d**, **f**, **h**) case-control GWAs SNPs. Var. *glauca* is depicted in purple (**a**, **b**, **c**, **d**), and var. *menziesii* in green (**e**, **f**, **g**, **h**). Background SNPs are sets of randomly sampled SNPs from their respective total set of SNPs for each variety and trait (drought tolerance and cold hardiness) after filtering out significant CMH SNPs (before FDR correction). The number of randomly sampled background SNPs for each set was the same as the number for significant SNPs. The significant SNPs were selected based on FDR corrected *p*-values from CMH tests (*q*-value ≤ 0.05), and on their overlap with significant (*p*-value ≤ 0.05) logisti regression SNPs (see Fig. 3 for the number of SNPs in each combination of trait and variety). Only one SNP per gene was considered for the heatmaps (lowest *q-value*). The average LD across all SNPs in each set 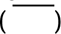 is depicted within each heatmap.

**Table 1.** Differentiation among populations for putatively neutral (background) and trait-associated SNPs (F_ST_), and phenotypes (v_POP_) within var. *glauca* (35 populations) and var. *menziesii* (39 populations), and between the two varieties. F_ST_ calculated with associated SNPs used the intersection between the significant SNPs from CMH test corrected for FDR (*q*-value ≤ 0.05) and the significant logistic regression SNPs (*p-value* ≤ 0.05). F_ST_ calculated with background SNPs used the average value from 100 random draws of non-significant SNPs of the same sample size as the associated SNPs. *p-values* in square brackets for associated SNPs’ F_ST_ values were obtained by testing for the number of trait-associated SNPs with significantly higher F_ST_ values than the 0.975 quantile of a null binomial distribution constructed with F_ST_ values from random SNPs drawn from the background. Values in square brackets after v_POP_ estimates are the standard errors of the estimates.

| Variety (trait) | $F_{ST}$ (associated SNPs) | $F_{ST}$ (background SNPs) | $V_{POP}$ (phenotype)* |
| --- | --- | --- | --- |
| Var. *glauca* (drought tolerance) | 0.043 [0.4985] | 0.031 | 0.140 [0.06] |
| Var. *glauca* (cold hardiness) | 0.071 [$<<0.001$] | 0.030 | 0.338 [0.10] |
| Var. *menziesii* (drought tolerance) | 0.042 [0.5022] | 0.026 | 0.001 [0.03] |
| Var. *menziesii* (cold hardiness) | 0.046 [0.0780] | 0.027 | 0.325 [0.10] |
| All 74 populations (drought tolerance) | 0.136 [0.0131] | 0.107 | 0.078 [0.03] |
| All 74 populations (cold hardiness) | 0.146 [0.0507] | 0.107 | 0.331 [0.07] |
| Var. *menziesii* vs. var. *glauca* (drought tolerance) | 0.156 [0.0039] | 0.121 | 0.250 [0.37] |
| Var. *menziesii* vs. var. *glauca* (cold hardiness) | 0.150 [0.0507] | 0.123 | 0.910 [1.26] |

\* $V_{POP}$ estimates for drought tolerance are in ref. 39, and for cold hardiness more details are in *SI Appendix*. $V_{POP}$ is a measure of genetic differentiation of phenotypic traits similar to $Q_{ST}$ (4).

Based on patterns for randomly chosen SNPs, we found genetic variation to be partitioned mostly within populations (*F_ST_* ≈ 0.03 for var. *glauca* and var. *menziesii*, Table 1), with about 3-fold stronger differentiation between var. *glauca* and var. *menziesii*. When the case-control candidate loci were used, *F_ST_* increased significantly for cold hardiness to *c.* 0.07 (*p-value* << 0.001), but not for drought tolerance *F_ST_* ≈ 0.04, *p-value* = 0.499; Table 1) was observed in var. *glauca*. Similarly, in var. *menziesii*, F estimate for cold hardiness associated loci increased to *c.* 0.05 (*p-value* = 0.078), but for drought tolerance, only to *c.* 0.04 (*p-value* = 0.502; Table 1). These results indicate a stronger signal of local adaptation to cold temperatures than drought in Douglas-fir, and partially confirm the weak signal of differentiation in the *V_pop_* estimate for drought tolerance within varieties (Table 1). Interestingly, the largest F observed was between the two varieties for drought-associated SNPs (F = 0.156), which was significantly larger than the background *F_ST_* (0.121, *p-value* = 0.004; Table 1), and suggests that the genetic architecture underlying drought tolerance in Douglas-fir is more differentiated between varieties than would be predicted by demographic history alone.

As highly polygenic local adaptation can result in elevated levels of LD (55), we next checked for differences in LD (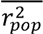) between putatively neutral (background) and trait-associated SNPs. In line with most *F_ST_* estimates, we observed a significant difference between LD in trait-associated SNPs and background LD, except for cold hardiness SNPs in var. *menziesii*. The average pairwise LD for drought tolerance and cold hardiness-associated SNPs in var. *glauca* (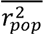= 0.064 and 0.065, respectively) was larger than the pairwise background LD (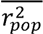 = 0.044 and 0.050, respectively; *p-value* < 0.001; Fig. 4a-d), with a less concentrated pattern of LD for cold hardiness-associated loci than the block of SNPs observed for drought tolerance-associated loci (Fig. 4b, d). In both cases, the potential concentrated effect of alleles in LD on phenotypes may have favored the detected signal of local adaptation for these traits in var. *glauca* (i.e., *V_pop_*; Table 1). The average pairwise LD for drought tolerance-associated SNPs in var. *menziesii* (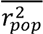 = 0.050) was significantly larger than the corresponding background LD (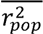 = 0.044; *p-value* < 0.001; Fig. 4e, f). The average pairwise LD for cold hardiness SNPs in var. *menziesii* was 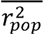 = 0.070, which did not differ from background LD (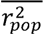 = 0.066; *p-value* = 0.39; Fig. 4g, h).

### Do loci with phenotypic associations also exhibit association with the environment?

If local adaptation is driving variation in drought tolerance and cold hardiness, we might expect to see spatial patterns of variation in allele frequency at loci strongly associated with these traits, covarying with the environmental variables driving local adaptation. To examine this, we tested the allele frequencies of the trait-associated SNPs–called for the 74 natural populations–for associations with 19 selected climate variables often hypothesized to be drivers of local adaptation in temperate conifers (33, 56) (*SI Appendix*, Table S1). To facilitate comparison between signals of phenotypic and environmental association with alleles, allele frequencies of highly associated case-control SNPs were transformed such that the frequencies (*p*) were converted to *1 – p* if the signal of the phenotypic effect was negative to ensure that all the studied alleles were positively associated with the traits (positive effect alleles - PEAs). We then used a multiple linear regression approach to regress the allele frequencies of the filtered PEAs on the first five principal components of a PCA with the 19 normalized climate variables. For each trait-associated SNP set, we selected the PEAs with the strongest relationship with climate (i.e., full-model adjusted R^2^ ≥ 0.4 and *p-value* ≤ 0.05). We then compared the number of PEAs highly associated with climate and their adjusted R^2^ with climate-associated SNPs obtained from random sets of SNPs across the genome (i.e., not associated with the traits) to test whether SNPs from the genomic background would produce similar patterns of association with the environment (see details in *SI Appendix*).

We found little difference in the strength of association to climate between phenotype-associated SNPs and randomly-chosen ones: the average adjusted-R^2^ for the association between PEAs and the first five climate principal components (PC) did not deviate from the null-distribution of adjusted-R^2^s for sets of random background SNPs from across the genome, correlated with the same PCs (Fig. 5a, c, e, g). Furthermore, the number of SNPs that passed the pre-determined threshold (adjusted-R^2^> 0.4) for the association with the climate PCs tended to be smaller for the phenotype-affecting SNPs than for the random background SNPs (Fig. 5b, d, f, h). Thus, while we did see evidence that both *F_ST_* and LD were often higher in SNPs associated with locally adapted phenotypes (Table 1; Fig. 4), alleles at these loci did not exhibit stronger linear associations with climate variables than those found randomly chosen loci.

**Fig. 5.**
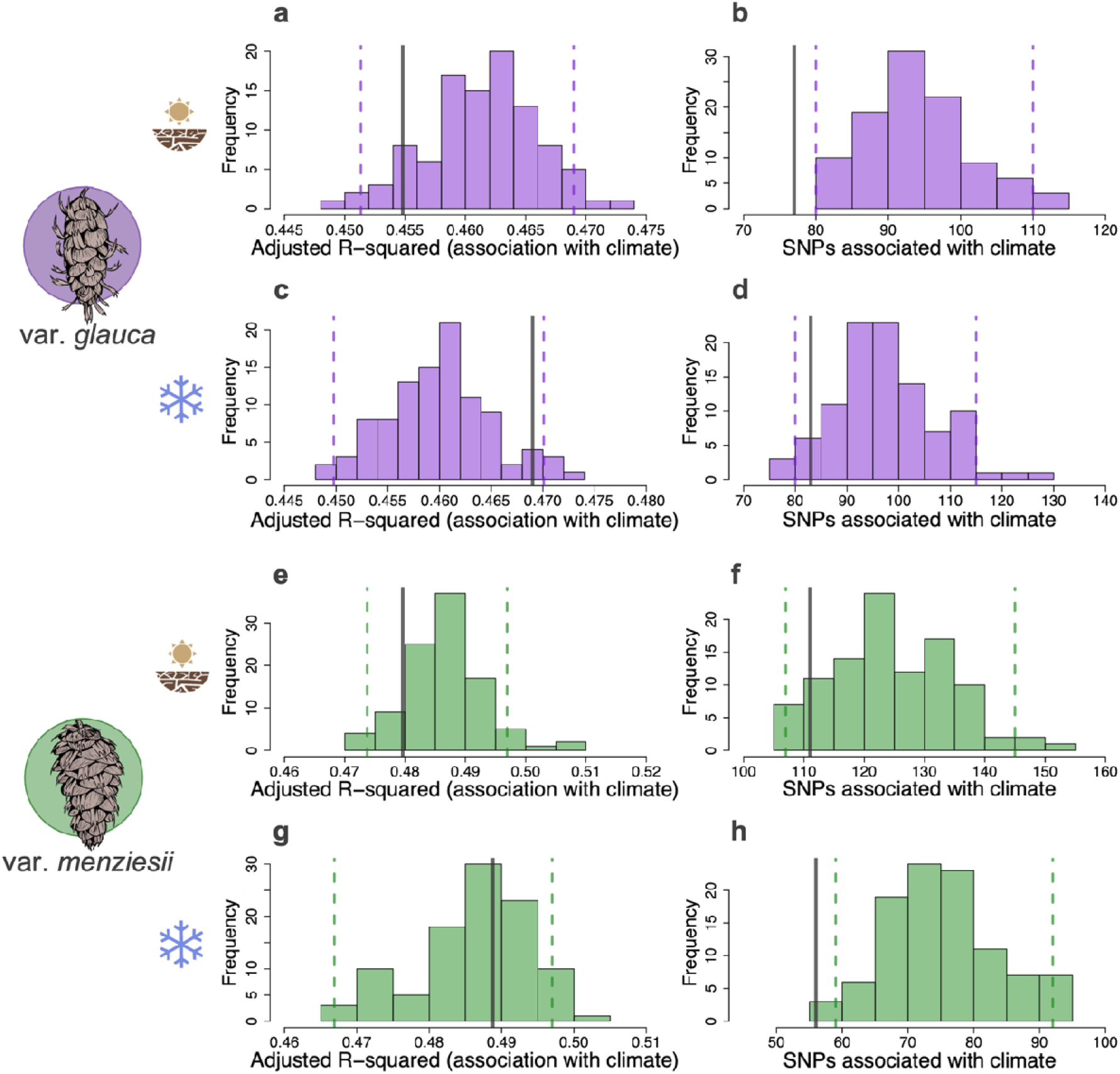
Patterns of association between the allele frequencies of random background SNPs and climate (histograms), and between positive effect alleles (PEA, i.e., from phenotype-affecting SNPs) and climate (gray vertical lines). **a**, **c**, **e**, **g**, depict the adjusted R^2^ distributions from the multiple linear regressions between the allele frequency of random background SNPs (100 independent draws from the genomic background with the same size of PEA SNP sets) and the first five principal components (PCs) of a climate PCA with 19 climate variables. **b**, **d**, **f**, **h**, depict the distribution of the number of random SNPs (from 100 independent draws from the genomic background with the same size of PEA SNP sets) that passed the same threshold used for PEA SNPs (adjusted R^2^ ≥ 0.4 and *p-value* ≤ 0.05). Gray lines depict the average adjusted-R^2^ and number of SNPs, highly associated with climate, from the multiple linear regressions between the PEA allele frequencies (i.e., SNPs associated with a trait) and the same first five principal components (PCs) of the climate PCA. Dashed lines depict the 0.025 and the 0.975 quantiles of their respective distributions (see details in *SI Appendix*).

To further characterize any spatial patterning of phenotype-associated alleles along climatic gradients, the climate PC-associated PEAs (i.e., associated with the five climate PCs) were first clustered using a Euclidean k-means algorithm as in ref. 26. The mean PEA frequencies of each cluster were then fitted against the populations’ environmental variables with a LOESS regression approach, and simple linear regressions. Phenotypic clines for cold hardiness along climatic gradients have been extensively reported in widespread temperate conifers (56–60), yet evidence for clinal variation in drought tolerance is limited (39) (but see refs. 61 and 62). Here, we found that variation in allele frequency of drought tolerance and cold hardiness PEAs was mostly driven by latitude and continentality (TD) in var. *glauca* (*SI Appendix*, Table S6, S7), and latitude and Hargreaves reference evaporation (Eref) in var. *menziesii* (Fig. 6b, *SI Appendix*, Table S8, S9). Nevertheless, while we detected clines in the frequency of PEAs for both traits and varieties, in general, we observed clusters of loci displaying both positive and negative associations between the allele frequencies and environmental variables (Fig. 6; *SI Appendix,* Fig. S8-13). This would not be expected based on extrapolation from a single-locus model of local adaptation (divergent directional selection), as in that case we would always expect the allele associated with an increase in tolerance to be found at higher frequency in the more stressful environment (e.g., allele associated with cold hardiness at higher frequency in populations with lower Mean Annual Temperature). Even for the traits that exhibit the strong phenotypic local adaptation, in fact, spatial patterning at the underlying loci occurs in conflicting directions, with some alleles that are positively associated with the trait showing negative associations with the environment and others showing the reverse. While it is difficult to make conclusive statements about the basis of local adaptation given the lack of difference in these patterns from background neutral loci, if anything, these patterns are suggestive of stabilizing selection towards different local optima in different regions, which can be driven by different combinations of alleles that increase and decrease the phenotype.

**Fig. 6.**
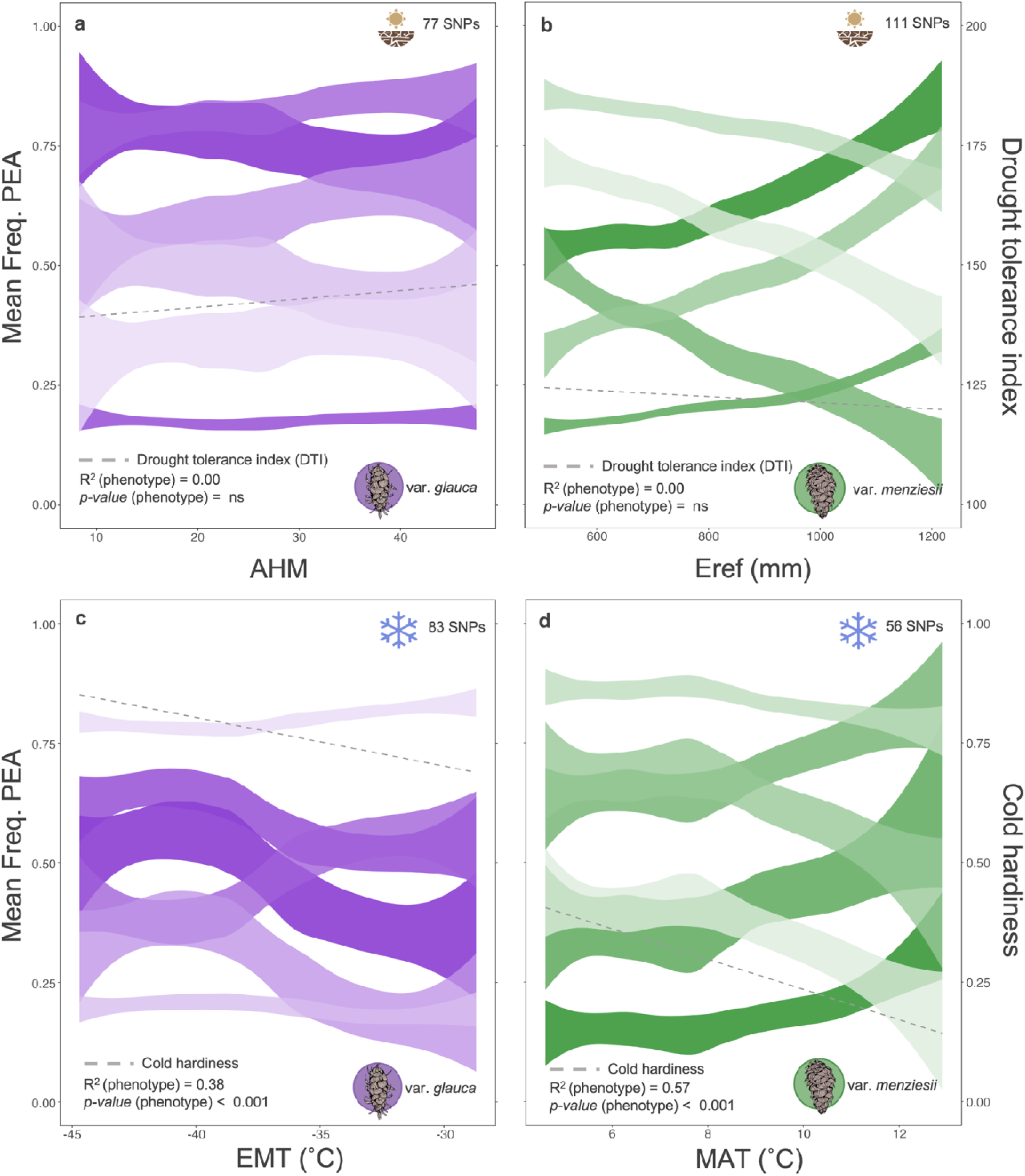
Genetic clines of phenotypes (gray dashed lines) and clustered (colored ribbons) drought tolerance (**a** and **b**) and cold hardiness (**c** and **d**) positive effect alleles along climate variables within var. *glauca* (**a** and **c**) and var. *menziesii* (**b** and **d**). The climate PC-associated PEAs for each trait and variet (i.e., each SNP set) are grouped into six clusters based on allele frequency similarities acros populations–35 for var. *glauca* and 39 for var. *menziesii*–spanning most of each variety’s natural range. The ribbons were generated with Loess regressions with a 0.8 span. The relationship between mean drought tolerance-associated PEA frequencies and annual heat-moisture index (AHM) for var. *glauca* and Hargreaves reference evaporation (Eref (mm)) for var. *menziesii* are depicted in **a** and **b**, respectively. The relationships between mean cold hardiness-associated PEA frequencies and extreme minimum temperature over 30 years (EMT (°C)) for var. *glauca* and mean annual temperature (MAT (°C)) for var. *menziesii* are depicted in **c** and **d**, respectively. Results for phenotypes (drought tolerance and cold hardiness; *SI Appendix*, Table S1) regressed on climate variables (simple linear regression) are depicted in each plot (i.e., R^2^ and *p-value*). See *SI Appendix* Fig. S8 and S9 for separate plots of each cluster, and Tables S6-9 for R^2^s and *p-values* of simple linear regressions between the mean PEA frequencies in each group and climate variables for the same populations.

As we did not observe consistent environmental associations at loci that were strongly associated with phenotypes, we searched for further evidence that they are involved in local adaptation by comparing the strength of phenotypic associations and environmental associations. To do this, we tested for a relationship between PEA effect sizes (log(Odds ratio)) and the slopes estimated from simple linear regressions of PEA frequencies regressed on environmental variables (under neutrality we would not expect an association between these effect sizes). The cold hardiness PEAs in var. *glauca* and drought tolerance PEAs in var. *menziesii* showed weak but significant relationships between the standardized slopes and the PEA effect sizes for most of the tested environmental variables (*SI Appendix*, Fig. S15, S16). Those significant relationships mostly agreed with the expectation that variables would drive stronger associations for alleles of larger effect (17). For instance, we observed more strongly positive slopes in the relationship of cold hardiness alleles with continentality (TD) for larger effect loci in var. *glauca* (R^2^ = 0.09; *p-value* = 0.006, *SI Appendix*, Fig. S15), and more strongly positive slopes for larger effect loci in the relationship between drought tolerance alleles and Hargreaves reference evaporation (Eref) in var. *menziesii* (R^2^ = 0.05; *p-value* = 0.021, *SI Appendix* Fig. S16). However, considering the large number of significant SNPs detected in our GWAS analyses (Fig. 2), and the small number of alleles that passed the minimum threshold of association with the multivariate environment used here (R^2^ ≥ 0.4; Fig. 6), along with the contrasting clines observed with most environmental variables, there are likely many loci affecting these traits and considerable redundancy.

## Discussion

With the growing pressure on forest ecosystems due to climate change, climate-adaptive traits such as drought tolerance and cold hardiness have become a major concern in breeding and management programs of temperate trees. Here we unveil key aspects of the genetic architecture of these two important traits in the two main varieties of *Pseudotsuga menziesii* (var. *menziesii* and *glauca*) and explore the distribution of alleles associated with geographic and climatic gradients across most of the species range. We found that both traits are highly polygenic in both varieties, with mostly low to moderate minor allele frequencies and small effect sizes (Fig. 2). Surprisingly, we observed little overlap in genes contributing to variation in drought tolerance and none in cold hardiness between the two varieties (Fig. 3), suggesting that different genetic architectures in each variety might be involved in the response to drought and freezing temperatures. In addition, drought tolerance and cold hardiness showed little or no pleiotropy, except for seven genes in var. *glauca* (Fig. 3), the variety with the strongest signals of local adaptation for both traits. Interestingly, drought tolerance, a trait with negligible phenotypic evidence for local adaptation within var. *menziesii* seedlings (Table 1), showed increased levels of LD for trait-associated SNPs, but with allelic clines along geographic and climatic gradients for different positive effect alleles (hereafter, simply alleles) in opposing directions (Fig. 6, *SI Appendix*, Fig. S12). Cold hardiness, on the other hand, a trait with moderate to strong local adaptation in Douglas-fir (Table 1; (41, 60, 63, 64), and other temperate conifers (4), exhibited moderate allelic clines, and more consistent relationships with temperature-related variables in var. *glauca* than var. *menziesii*, but still not in full agreement with that expected under simple divergent directional selection (65). Altogether, our results suggest that most of the genomic variation underlying these two important climate-adaptive traits is remarkably complex, with genetic redundancy likely playing a large role in the evolution of local adaptation in the two varieties. This has significant implications for genomic predictions of maladaptation to climate change, for informing management and conservation, as well as for breeding climate-adapted trees–linear clines along environmental axes can be observed at loci that affect the phenotype, but, many times, in the direction opposite that which would be predicted based on the match between phenotype and environment. Thus, there would be a mismatch between a model predicting response to climate change based on a per-locus phenotypic effect vs. a per-locus environmental effect (i.e., the approach commonly used with genomic offsets).

Trait architectures with few large-effect loci, often concentrated in genomic regions, might be favored by local adaptation over time if the fitness optimum varies in space and selection is stronger than migration, but if traits have a highly polygenic basis with redundancy, such predictions may not be realized, particularly if there is also temporal variability in the environment (17, 66). At the scale of variation among populations within varieties and within the entire species, we observed a substantially smaller signal of local adaptation to drought than to cold temperatures at the phenotypic and genomic levels (Table 1). Our results suggest that local adaptation in Douglas-fir could be primarily driven by low temperatures in both varieties, while drought tolerance contributes more weakly to local adaptation in var. *glauca* (Table 1), and even less so in var. *menziesii*. These contrasting results also suggest that, despite an apparent abundance of genetic variation segregating within-populations for both traits (Fig. 3), drought may have been a less efficient selection pressure than freezing temperatures in the Pacific Northwest, potentially due to higher spatial and temporal variability (i.e., lower spatiotemporal autocorrelation), or the unpredictability of droughts (as discussed in ref. 39). Spatial autocorrelation of a selection pressure can have significant impact on patterns of local adaptation (24), but the connection between temporal autocorrelation and patterns of local adaptation still needs further investigation (17).

It was previously suggested that gene flow may swamp drought-associated alleles and prevent local adaptation to drought as evidenced by phenotypic variation in var. *menziesii* (39). The relatively continuous distribution of populations in the PCA analysis of var. *menziesii* (*SI Appendix*, Fig. S7e, g), combined with low *F_ST_* estimates for background SNPs in both varieties, indicate strong gene flow. High levels of long-distance gene flow are widespread among temperate conifers (67). However, the large standing genetic variation maintained within populations for drought tolerance at the phenotypic and genomic level and the lack of evidence of local adaptation for phenotypes, could also suggest that drought tolerance may be governed by a genetic architecture with redundant loci, or the trait might have evolved through global rather than local adaptation (28), in particular within var. *menziesii*. Given the increase in the frequency and intensity of droughts in the region with climate change and the observed standing genetic variation, over time, greater adaptation to drought could still evolve (13).

Average LD across all loci associated with both traits in var. *glauca* was significantly higher than the background LD (*p-value* << 0.001, Fig. 4a, c). LD can maintain phenotypic differentiation, even for polygenic traits (28). When multiple associated loci are tightly linked, they can segregate as a supergene (*sensu* ref. 68), upon which selection can act more effectively, maintaining variation. Large LD blocks were previously identified for genotype-environment associated (GEA) loci in var. *glauca*, particularly those associated with precipitation related variables (53), and we observed some clustering among loci that was suggestive of this (e.g., Fig. 4b). However, we do not have a linkage map and cannot confirm whether SNPs in LD clusters are physically linked or distributed across chromosomes. Among-chromosome LD might not be strong enough to overcome the effects of migration and recombination (28). On the other hand, genomic regions in high LD can also be caused by structural variations (e.g., such as inversions), reducing recombination and facilitating local adaptation (69–71). Altogether, LD patterns in Douglas-fir corroborate our other results showing that climate-associated traits are more locally adapted in var. *glauca* than in var. *menziesii*.

The limited overlap between varieties for genes underlying drought tolerance (five genes, *p-value* = 0.052; Fig. 3; *SI Appendix*, Table S3), and the lack of overlap in cold hardiness genes (*p-value* = 0.31), suggests that the two varieties may have evolved different biochemical, physiological, morphological or phenological adaptations or been using variation in different genes to deal with drought and cold (72–74). The traits we used to assess drought tolerance (rate of decline in photosynthetic efficiency) and cold hardiness (proportion of electrolyte leakage after freezing) are proxies for survival under these stresses, and can be considered high-level phenotypes, at the top of the trait hierarchy (*sensu* ref. 75) Alternative lower-level phenotypes, such as gene regulatory mechanisms, may yield similar higher-level phenotypic outcomes. For instance, var. *menziesii* seedlings tend to experience an earlier decline in photosynthetic efficiency under drought treatment (39), likely caused by stomata remaining open (i.e., anisohydric strategy), which can ultimately result in hydraulic failure (76). Meanwhile, var. *glauca* seedlings are only likely to reduce photosynthetic efficiency at much later stages (39), indicating an avoidance strategy with stomata remaining closed to maintain a stable water potential (i.e., isohydric strategy), which could result in death from carbon limitations (76). These two strategies may involve different stress signaling mechanisms (77), evidenced by the low repeated evolution observed here, and which in turn, will induce different higher-level phenotypes. These, however, can eventually result in similarly tolerant trees or populations between the two varieties (39). Likewise, different metabolic mechanisms controlling cold acclimation were observed among distant populations of *Picea sitchensis*, suggesting different cold hardiness activation strategies within the same species (78). Future studies of repeated climate adaption between var. *glauca* and var. *menziesii* could tackle traits from multiple levels of the trait hierarchy, increasing the chances of detecting repeatability.

A complementary explanation for the low level or lack of repeated evolution between the two varieties relates to how drought tolerance and cold hardiness correlate with other adaptive traits, potentially more strongly affected by selection for local adaptation in each variety (e.g., growth competition in var. *menziesii*, 79). For example, early tolerance to drought in seedlings has an antagonistic correlation with height growth among populations of var. *menziesii* (39). In this case, selection for height growth might counteract development of optimal phenotypes for early drought, as the fitness cost to be drought tolerant may be too high when competition is also high. Meanwhile, var. *glauca* seedlings show a moderate positive, reinforcing correlation between drought tolerance and height growth among populations (39). This suggests that at the phenotypic level in var. *glauca*, these two traits are coupled and respond to selection in concordant directions, as to survive and grow in colder and drier continental climate conditions, trees need to tolerate more abiotic stress than var. *menziesii* in maritime climates. Repeated evolution has been found to be higher for traits under more consistent selection pressures (31), with stronger pleiotropy among traits if they contribute to local adaptation (36), or with weaker pleiotropy if they mostly contribute to global adaptation (52). The extent of local adaptation of drought tolerance in the two varieties reported here (V = 0.01 (var. *menziesii*), and 0.14 (var. *glauca*), Table 1), and of height growth increment reported in ref. 39 (*V_pop_* for height increment = 0.22 (var. *menziesii*), and 0.06 (var. *glauca*)) further supports this argument. The taxonomic proximity between the two Douglas-fir varieties (divergence time *c.* 2.1 million years (43)) does not seem to be an important factor contributing to repeatability in the species, in contrast with general expectations (34).

In conifers, the most severe physiological impacts caused by freezing and drought events can be similar, such as hydraulic failure and photosynthetic impairment (8). Consistent with this, a positive covariation between cold and drought tolerance was previously observed in colder provenances of var. *menziesii* (80), indicating the potential for pleiotropy. Surprisingly, we did not detect a single gene associated with both cold and drought tolerance in var. *menziesii* (hypergeometric test *p-value* = 0.29) and found only seven overlapping genes in var. *glauca* (*p-value* = 0.01; Fig. 3). The larger number of pleiotropic genes observed in var. *glauca* relative to the total number of significant genes detected for both traits suggests these two traits evolved less independently in var. *glauca* than in *menziesii*. The colder and drier environments where *glauca* occurs may have more predictable and consistent selection pressures than in the generally warmer and wetter, highly productive coastal environment of var. *menziesii*, where competition for growth may be stronger (79). Additionally, despite the expectation for plants to use some of the same gene regulatory mechanisms to respond to these two stressors (38), sensing and signaling pathways activating responses to either freezing temperatures or drought appear markedly independent in our study. In general, while much of the response to freezing in conifers is triggered by Ca^2+^ sensing and signaling pathways (73), responses to drought are largely related to the production and signaling of abscisic acid (ABA) (38, 72). In fact, important genes related to Ca^2+^ signaling and response to freezing, such as CIPK20 (81) and UBC13 (82), and genes important for ABA signaling and drought tolerance, such as CHLH (83) and CKX5 (84) were identified in our annotations for cold hardiness-associated and drought tolerance-associated SNPs, respectively, but no annotated higher-level response gene overlapped between traits.

Pleiotropy may be enriched in traits that contribute to local adaptation to climate, which, in turn, may increase chances of convergence among plant species (36). This is consistent with the stronger signals of local adaptation and overlapping genes we observed between drought tolerance and cold hardiness in var. *glauca* when compared to var. *menziesii* (Fig. 3b). Given the coupling of cold hardiness and growth traits in Douglas-fir (60) and the well documented local adaptation for growth traits (41, 42, 60, 85), future studies should assess the extent of pleiotropy between climate-adaptive traits and growth traits in Douglas-fir. Such studies could identify genes involved in trade-offs between growth and climate-adaptive traits, and inform predictions of evolutionary responses to climate change.

Surprisingly, latitude rather than a climatic variable was most strongly associated with drought tolerance and cold hardiness loci, despite the lack of associations between latitude and these traits in var. *glauca* and only a weak association between latitude and cold hardiness in var. *menziesii* (*SI Appendix*, Table S4, S5). Latitude can be considered a proxy for some other environmental variables such as temperature and photoperiod, which are typically associated with variation in phenological traits and are drivers of local adaptation in many widespread trees (4, 66), but latitude does not capture the steep climatic gradients from maritime to continental climates. The strong latitudinal variation in allele frequencies could be the result of variation in phenological traits genetically correlated with cold and drought tolerance. Alternatively, postglacial expansion and colonization could also have generated such latitudinal clines through allele surfing (86). Var. *menziesii* most likely had a single refugium during the Last Glacial Maximum from which northward expansion took place (43), potentially reflecting the observed sharp latitudinal clines in allele frequencies within this variety. In contrast, var. *glauca* likely had multiple refugia, but the colonization routes leading to our sampled populations are less clear (43).

Whether or not local adaptation could evolve without allelic clines in underlying loci across environmental gradients is an important question, as it challenges one of the main assumptions of GEA methods (25). GEAs only detect significant relationships; they do not assess the direction of relationships between alleles and environments, nor can they determine if alleles are associated with phenotypes. In addition, if a trait is controlled by multiple loci of small effects, redundancy is expected (27). Loci important for adaptation in one population might not be the same as those in other populations, which may generate variation in allele frequencies among populations that is not associated with environment. This leads to more complex questions when using genomic data to predict adaptation to climate. For example, to what extent will genetic redundancy affect predictions of future maladaptation based on genomic data? Even if clines exist for alleles underlying adaptive traits, would alleles vary in effect size among populations? The lack of a consistent pattern in var. *menziesii* cold hardiness allele frequencies along geographic and climate variables, for example, suggests that redundancy and varying allele effect sizes among populations may both be occurring (*SI Appendix*, Fig. S13, S17). Despite the moderate phenotypic clines along environmental gradients, some allele clusters agreed with the variation in the same direction along the same gradients, while others were in the opposite direction (*SI Appendix*, Fig. S13). In this case, GEA methods would not have been appropriate for studying in detail the genetic architecture of local adaptation to cold temperatures. Correctly predicting maladaptation to future climates with genomic offset methods based on GEAs would also be challenging in this case, as the predicted future genomic composition may not reflect actual phenotypic adaptive response. Recent studies in temperate conifers have suggested that genomic offset predictions using GEA-identified loci often perform similarly to randomly selected putatively neutral loci across the genome (87, 88), and this could be another indication of the redundant architectures controlling climate-adaptive traits (29).

### Limitations

The selection and pooling of individuals from the extremes of the phenotypic distribution in each population before performing the GWAS analyses is a limitation in our study. The grouping of continuously distributed drought tolerance and cold hardiness phenotypes into cases and controls may have reduced our ability to detect additional small-effect loci and precluded the use of haplotype information. This experimental design also restricted our analysis to GWAS approaches that support binary data only (e.g., CMH test and logistic regression) and did not down-weight contributions from populations with minimal phenotypic differences between case and control. On the other hand, the pooling of individuals allowed us to assess a much larger phenotypic variation for the same number of genotyped samples given the number of assessed populations and that the extreme pools were selected from up to 95 individuals in each population. We did not have access to a chromosome-level reference genome for either variety, and the mapping of var. *glauca*’s SNPs was done onto the available reference genome for var. *menziesii,* limiting our ability to detect more variants and interpret signatures in light of their linkage relationships.

### Conclusion

The interaction between traits, the specificity of loci within varieties and possibly within populations, and the strong indication of genetic redundancy underlying drought tolerance and cold hardiness is evidence of remarkably complex polygenic architectures controlling for climate adaptation in Douglas-fir. This complexity currently limits the ability of using solely genomic information to inform climate-adaptive management in Douglas-fir.

These results add to concern over the use of GEA loci in genomic offset studies to characterize and manage species for climate-adaptive traits with complex polygenic architectures. Using the putative drought tolerance adaptive loci identified in this study to predict maladaptation to future drought conditions is not recommended. On the other hand, the agreement observed between phenotypic and allelic clines in cold hardiness, particularly within var. *glauca*, suggests cold hardiness-associated loci could provide a good indication of the thermal tolerance of populations and the extent of mismatch with future climate conditions based on genomic offset predictions. Given such diffuse genomic architectures for these two climate-adaptive traits in seedlings of Douglas-fir, management decisions may still be better informed by well-established common garden experiments and well-defined and assessed phenotypes.

## Materials and Methods

### Sampling

We selected 20 Douglas-fir (*Pseudotsuga menziesii* (Mirbel) Franco) wild populations spanning most of the natural range of var. *menziesii* (11 populations) and var. *glauca* (9 populations) (Fig. 1a) to assess within-population variation and identify genes associated with drought tolerance and cold hardiness in two separate common garden experiments (Fig. 1c, d). Open-pollinated seedlots, collected from multiple parents for operational reforestation, were obtained for each population from existing collections in seedbanks in the USA and Canada (https://coadaptree.forestry.ubc.ca/seed-contributors/). Var. *menziesii* provenances span more than 11° latitude, 6° longitude (∼490 km), and 1250 m elevation, and var. *glauca* provenances more than 8° latitude, 11° longitude (∼760 km), and 1000 m elevation, representing a large range of environmental conditions (*SI Appendix,* Fig. S1, Table S1).

### Drought tolerance phenotyping and selection of cases and controls

The drought experiment comprised 74 to 80 seedlings per population submitted to a drought-to-death treatment in a greenhouse between April 2018 and October 2018 in Vancouver, BC, Canada (49.25°N, 123.23°W). One-year old seedlings from each population were transplanted before bud flush into a single large, multi-seedling box (13□×□35□×□90□cm) made from plastic core board (Fig. 1c), with 20 boxes total (i.e., one per population). The central 80 seedlings in each box were measured, totaling 1,594 measured seedlings. The drought treatment started after eight weeks of acclimation, during which field capacity was maintained in all boxes. After two weeks of gradual water reduction (relative to the field capacity), watering was completely halted for two weeks. A volumetric water content (VWC) of *c.* 5% was maintained in all boxes until the end of the experiment. See *SI Appendix* for full details of the experiment.

The decline in photosynthetic efficiency of all seedlings under drought was assessed based on six measurements of chlorophyll fluorescence (F_v_/F_m_(89)) during the time of the experiment. Potential effects of spatial autocorrelation of drought on the measurements in each box were removed with an autoregressive model of residuals implemented in ASReml-R 4.0 (39, 90). The drought tolerance index (DTI) was then estimated as the random slope of a mixed effect model implemented in the NLME package in R (91) for each population (i.e., box), where the corrected F_v_/F_m_ was the dependent variable and the interaction between days of the experiment and seedling ID was considered a random effect (*SI Appendix*). Therefore, DTI represents the average decline in photosynthetic efficiency over time due to drought. The ten least (cases) and the ten most (controls) drought tolerant individuals were then selected within each population (*SI Appendix,* Fig. S2), based on which case-control GWAS was subsequently performed.

### Cold hardiness phenotyping and selection of cases and controls

The cold-hardiness experiment was established in outdoor nursery raised beds in March 2018 in Vancouver, BC, Canada (49.26°N, 123.25°W). Approximately one year after germination and before bud flush, between 78 and 95 seedlings per population, from the same 20 populations, were transplanted into a randomized unbalanced block design experiment with 11 blocks (Fig. 1d), totaling 1,839 seedlings being tested for cases and controls. Seedlings from 54 additional natural populations were included in the cold hardiness experiment (12 seedlings on average per population) for the assessment of phenotypic differentiation and clines within varieties (see Fig. 4i and *SI Appendix*, Table S1 for more details). We assessed fall cold-injury (%) based on electrolytic leakage of detached needles after artificial freezing (92). See *SI Appendix* for full details of the experiment and measurements. The effect of block, population, and conductivity meter was removed from the cold injury values with a beta regression approach (93) implemented in the BETAREG package in R (94), ideal for analyzing proportions derived from continuous data (95). Different combinations of variables and interactions was tested and the model with the lowest AIC was chosen for each tested temperature and variety (*SI Appendix*). The residuals of the selected model were then used as the corrected cold injury for each seedling. Ten cases (least hardy seedlings) and ten controls (most hardy seedlings) were then selected per population for the subsequent case-control GWAS analyses (*SI Appendix*, Fig. S3).

### Targeted exome capture pool-sequencing and SNP calling

Needles from all seedlings in the two experiments (drought and cold) were collected and stored in silica prior to the phenotypic measurements. We extracted the DNA from each of the 20 (ten cases and ten controls) phenotypically selected seedlings per population in each of the experiments separately. DNA pools of cases and controls for each population and trait were established with normalized contributions of DNA (same volume and concentration) from each individual (96). In total, 80 DNA pools were established (20 populations x two traits x (1 case + 1 control pool)), and pool-sequenced with a 50 Mb targeted sequence capture panel focused primarily on exonic regions (further details in *SI Appendix*). Each library (pool) was sequenced on an Illumina NovaSeq 6000 System S4 (PE150bp) by the Genome Quebec Innovation Centre (McGill University, Montreal, QC, Canada).

We used a pool-seq SNP calling bioinformatics pipeline (44), described in detail in ref. 100, to process our genomic data. Each of four datasets, one for each variety-trait combination, was processed independently. For further details on the SNP calling procedures, see *SI Appendix*. After calling SNPs, we calculated the average global allele frequency within pools for each Douglas-fir variety and trait by summing the products of the ALT allele frequencies and the ploidy of the pools (2N = 20), and then dividing it by the total ploidy across populations. We then filtered the SNPs for global minor allele frequency (MAF) ≥ 0.05.

### Case-control GWAS approaches

After transforming the assessed phenotypes into binary data separating cases (least tolerant) and controls (most tolerant), we performed two case-control GWAS analyses for each dataset: a) a modified Cochran–Mantel–Haenszel test (CMH) implemented with a PYTHON script (45), with correction for false discovery rate (FDR); and b) a logistic regression model implemented with a GLM function in R *4.2.0* (98), following recommendations by ref. 99. We considered only the significant SNPs from the intersection of these two approaches for downstream analyses. See *SI Appendix* for further details on the case-control GWAS approaches. As the significant SNPs from the logistic regression approach were solely used to validate the FDR-corrected significant SNPs from the CMH test, we did not correct the logistic regression p-values for false discovery rate to reduce false negatives. The *p-value* threshold used for the logistic regression SNPs was 0.05.

### Annotation and overlap of genes between varieties and traits

Transcriptomic contigs (i.e., genes) were identified from the reference Douglas-fir genome annotation (version 1.0; ref. 100) and from additional transcriptomic studies (49, 97). For details on the assembly, mapping, and annotation of the additional transcripts, see ref. 49. The significance of overlapping genes between the two varieties and between traits was tested with the hypergeometric test in R *4.2.0* (98). Finally, we performed Gene Ontology (GO) enrichment analysis of genes using TOPGO (version 2.56.0; 101).

### Variation of drought tolerance and cold hardiness associated alleles across the species range

SNPs were also called for the additional dataset of 74 natural Douglas-fir populations (39 var. *menziesii* and 35 var. *glauca*) from across the species natural range (*SI Appendix*, Fig. S1 and Table S1), which included the 20 populations used for the case-control GWAS but with an independent sample of seedlings (53). Seedlings from these 74 natural populations were grown in a separate drought experiment (see ref. 39) and in the same case-control cold hardiness experiment and were pool-sequenced with the same targeted exon captures used for our cases and controls, with each pool including 33 to 40 randomly selected individuals from one population (see ref. 53 for sequencing and SNP calling procedures for these 74 natural populations). Frequencies of SNPs associated with drought tolerance and cold hardiness for these 74 natural populations were used for subsequent analyses.

### Genetic differentiation and linkage disequilibrium

After calling the SNPs for the 74 natural populations, we used principal component analysis (PCA) to visualize potential structure among populations depending on the SNP set used in the analysis (i.e., a random sample of SNPs from the background, and the significant set of SNPs after FDR correction for each trait and variety), using the function *prcomp* in the package STATS in R *4.2.0* (98; *SI Appendix*). We also estimated *F_ST_* values among populations within varieties, species-wide, and between the two varieties for each set of top trait-associated SNPs (FDR corrected), and for random samples of background (non-trait-associated) SNPs with the package POOLFSTAF (version 2.2.0; ref. 102) in R *4.2.0* (98). To test the significance of the F values for trait-associated SNPs we checked for the number of SNPs that had an *F_ST_* higher than the 0.975 quantile of a null distribution of *F_ST_* values for random non-significant SNPs drawn from the background with the same sample size as the trait-associated SNPs. This process was repeated 100 times for each trait and variety dataset and the average number of SNPs above the 0.975 quantile of the null distribution across draws was obtained for each dataset. A binomial test was then conducted in R *4.2.0* (98) to test whether the average number of associated SNPs above the 0.975 quantile cut-off was higher than the expected by chance.

We also reported the phenotypic differentiation (*V_pop_*) for each trait and their respective standard errors within and between the two varieties, and species-wide, following the approach used in ref. 54. For this we used the cold hardiness phenotypes assessed for the same 74 natural populations in the cold hardiness experiment and the drought tolerance phenotypes previously reported in ref. 39. Next, we estimated a proxy for pairwise linkage disequilibrium (LD) among SNPs within each dataset for associated and non-associated SNPs. LD was estimated as the squared correlation between the frequencies of alleles across all natural populations within varieties (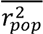). The difference between the estimated r^2^ for background SNPs and the associated SNPs was performed with the non-parametric test Wilcox signed-rank test in R *4.2.0* (98).

### Allelic clines across climate space

Climate normals (1961–1990) of 19 selected climate variables–often used in studies of local adaptation in temperate conifers (33, 56) were estimated for each provenance with ClimateWNA (103) (see list of variables in *SI appendix*, Table S1). To estimate allelic clines along climate gradients, we used a similar approach to ref. 26. First, we further filtered the SNPs for MAF ≥ 0.1 and only used the SNP with the lowest CMH *p-value* amongst the significant SNPs in each gene. For each trait-associated SNP set, we selected the SNPs with the strongest relationship with climate (full-model adjusted R^2^ ≥ 0.4 and *p-value* ≤ 0.05) based on a multiple linear regression approach, where the allele frequencies of the trait-associated SNPs were regressed on the first five principal components of a PCA with the 19 normalized climate variables. The selected climate-associated SNPs were compared to a null distribution constructed with randomly selected SNPs from the genomic background to test whether the patterns of association of trait-affecting loci with environment (i.e., adjusted-R^2^s and number of SNPs) were, or were not, significantly different than randomly chosen alleles from the genomic background (see details in *SI Appendix*, and Fig. 5).

Based on the observed effect of each trait-associated SNP on the traits in the GWAS analysis (logistic regression), we converted the allele frequencies (*p*) into *1 – p* if the signal of the effect was negative to ensure that all the selected alleles were positively associated with the traits (positive effect alleles–PEAs, ref. 50). The selected PEAs for each trait and variety were then clustered into six groups with similar frequencies across populations, with a Euclidian k-means algorithm in the package STATS in R *4.2.0* (98) The mean PEA frequency of each cluster was calculated for each population and plotted against each environmental variable of interest for each trait with a local estimated scatterplot smoothing (LOESS) approach. We used this non-parametric approach to look for potential localized clines along the range of the two varieties, which would not be apparent with simple linear regressions (26). We also fitted simple linear regressions to test for linear relationships between the mean PEA frequencies of each cluster across populations and their environmental gradients. Additionally, for comparison with the PEA clines, we present phenotypic clines for drought tolerance (previously reported in ref. 39), and for cold hardiness (*SI Appendix*, Table S1) across the 74 natural populations (i.e., the additional dataset) along climatic gradients for each variety.

Finally, in order to infer the potential for genetic redundancy among the associated SNPs within each trait, we looked for significant relationships between standardized slopes estimated from simple linear regressions between single PEA frequencies and the environmental variables, and the PEA effect sizes (log(Odds ratio)). The slopes were standardized by the mean PEA frequency, given that intermediate frequencies will produce higher Delta-p. The premise of this analysis is that, if a significant relationship is observed between the PEA standardized slopes and their effect sizes (i.e., there is a tendency of larger effect loci to drive stronger allelic clines along the environment for a locally adapted trait), the genetic redundancy of the trait is sufficiently low to allow at least some of the largest alleles to experience sufficiently strong selection to accentuate their divergence and contribution to local adaptation. On the contrary, if no consistent relationship is observed, there is potential evidence for genetic redundancy. It is important to emphasize that only SNPs that showed a strong relationship with the multivariate environment in the first place were used for this analysis (the same loci used in the PEA clines analysis).

## Data availability

Datasets will be made available on Dryad upon acceptance of this paper and before publication.

## Acknowledgements

This research was part of the CoAdapTree project led by S.N.A. and S.Y. and funded by Genome Canada (241REF; Co-Project Leader Richard Hamelin), with co-funding from Genome BC and sponsors including Genome Alberta, Génome Québec, the British Columbia Ministry of Forests, Canadian Forest Service (Natural Resources Canada), Alberta Innovates Bio Solutions, Vernon Seed Orchard Co., University of Alberta, University of British Columbia, the Forest Genetics Council of British Columbia, Digital Research Alliance of Canada, Mosaic Forest Management, TimberWest, and Western Forest Products. Seeds were donated by 16 forest companies and agencies in Canada, United States, and Mexico (for details, visit https://coadaptree.forestry.ubc.ca/seed-contributors/). We thank Rebecca Jordan, Stephen R. Keller, Hayley Tumas, and Rob Guy for their helpful feedback on the manuscript. We also thank Pia Smets, Tongli Wang, Colin Mahony, Yue Yu, Alex Girard, Iain Reid, Justin Chow, Beth Roskilly, Jon Degner, Susannah Tysor, and Joanne Tuytel for phenotyping assistance, and the Aitken Lab for helpful discussions. Eduardo Cappa, Ian MacLachlan, and Jon Degner provided helpful suggestions on the analyses. Martin Henry made the Douglas-fir cone drawings used in the figures. Fig. 1-6 were partially created in *BioRender*. Candido-Ribeiro, R. (2025) (https://BioRender.com/x67a953).

## References

1. P. K. Thornton, P. J. Ericksen, M. Herrero, A. J. Challinor, Climate variability and vulnerability to climate change: A review. Glob Chang Biol 20, 3313–3328 (2014).

2. J. Wang, N. B. Grimm, S. P. Lawler, X. Dong, Changing climate and reorganized species interactions modify community responses to climate variability. Proceedings of the National Academy of Sciences 120, e2218501120 (2023).

3. S. N. Aitken, S. Yeaman, J. A. Holliday, T. Wang, S. Curtis-McLane, Adaptation, migration or extirpation: climate change outcomes for tree populations. Evol Appl 1, 95–111 (2008).

4. F. J. Alberto, et al., Potential for evolutionary responses to climate change - evidence from tree populations. Glob Chang Biol 19, 1645–1661 (2013).

5. L. Leites, M. Benito Garzón, Forest tree species adaptation to climate across biomes: Building on the legacy of ecological genetics to anticipate responses to climate change. Glob Chang Biol (2023).

6. S. Zhou, Y. Zhang, A. P. Williams, P. Gentine, Projected increases in intensity, frequency, and terrestrial carbon costs of compound drought and aridity events. Sci Adv 5, 1–9 (2019).

7. J. Cohen, L. Agel, M. Barlow, C. I. Garfinkel, I. White, Linking Arctic variability and change with extreme winter weather in the United States. Science (1979) 373, 1116–1121 (2021).

8. K. A. McCulloh, et al., At least it is a dry cold: the global distribution of freeze-thaw and drought stress and the traits that may impart poly-tolerance in conifers. Tree Physiol 43, 1–15 (2023).

9. J. T. Anderson, The consequences of winter climate change for plant performance. Am J Bot 110, 0–2 (2023).

10. E. Kirchhof, F. Campos-Arguedas, N. S. Arias, A. P. Kovaleski, Thresholds for spring freeze: measuring risk to improve predictions in a warming world. New Phytologist 248, 563–575 (2025).

11. D. G. Bock, et al., Genomics of plant speciation. Plant Commun 4 (2023).

12. R. D. H. Barrett, D. Schluter, Adaptation from standing genetic variation. Trends Ecol Evol 23, 38–44 (2008).

13. D. N. Anstett, et al., Evolutionary rescue during extreme drought. bioRxiv 1–15 (2024). 10.1101/2024.10.24.619808.

14. K. Bomblies, C. L. Peichel, Genetics of adaptation. Proc Natl Acad Sci U S A 119 (2022).

15. S. N. Aitken, R. Jordan, H. R. Tumas, Conserving Evolutionary Potential: Combining Landscape Genomics with Established Methods to Inform Plant Conservation. Annu Rev Plant Biol 75, 707–736 (2024).

16. N. Isabel, J. A. Holliday, S. N. Aitken, Forest genomics: Advancing climate adaptation, forest health, productivity, and conservation. Evol Appl 13, 3–10 (2020).

17. S. Yeaman, Evolution of polygenic traits under global vs local adaptation. Genetics 220 (2022).

18. C. Rellstab, F. Gugerli, A. J. Eckert, A. M. Hancock, R. Holderegger, A practical guide to environmental association analysis in landscape genomics. Mol Ecol 24, 4348–4370 (2015).

19. S. Lachmuth, T. Capblancq, A. Prakash, S. R. Keller, M. C. Fitzpatrick, Novel genomic offset metrics integrate local adaptation into habitat suitability forecasts and inform assisted migration (2024).

20. D. Lazic, et al., Genomic variation of European beech reveals signals of local adaptation despite high levels of phenotypic plasticity. Nat Commun 15, 8553 (2024).

21. T. Capblancq, M. C. Fitzpatrick, R. A. Bay, M. Exposito-Alonso, S. R. Keller, Genomic Prediction of (Mal)Adaptation across Current and Future Climatic Landscapes. Annu Rev Ecol Evol Syst 51, 245–269 (2020).

22. J. R. Lasky, E. B. Josephs, G. P. Morris, Genotype-environment associations to reveal the molecular basis of environmental adaptation. Plant Cell 35, 125–138 (2023).

23. A. M. Waldvogel, et al., Evolutionary genomics can improve prediction of species’ responses to climate change. Evol Lett 4, 4–18 (2020).

24. T. R. Booker, The structure of the environment influences the patterns and genetics of local adaptation. Evol Lett 1–12 (2024). 10.1093/evlett/qrae033.

25. K. E. Lotterhos, The paradox of adaptive trait clines with nonclinal patterns in the underlying genes. Proceedings of the National Academy of Sciences 120, e2220313120 (2023).

26. C. R. Mahony, et al., Evaluating genomic data for management of local adaptation in a changing climate: A lodgepole pine case study. Evol Appl 13, 116–131 (2020).

27. Á. J. Láruson, S. Yeaman, K. E. Lotterhos, The Importance of Genetic Redundancy in Evolution. Trends Ecol Evol 35, 809–822 (2020).

28. S. Yeaman, Local adaptation by alleles of small effect. American Naturalist 186, S74–S89 (2015).

29. K. E. Lotterhos, The paradox of adaptive trait clines with nonclinal patterns in the underlying genes. Proceedings of the National Academy of Sciences 120, e2220313120 (2023).

30. L. Wang, et al., Molecular Parallelism Underlies Convergent Highland Adaptation of Maize Landraces. Mol Biol Evol 38, 3567–3580 (2021).

31. S. Soudi, et al., Repeatability of adaptation in sunflowers reveals that genomic regions harbouring inversions also drive adaptation in species lacking an inversion. Elife 12, 1–38 (2023).

32. M. Bohutínská, et al., Genomic basis of parallel adaptation varies with divergence in Arabidopsis and its relatives. Proc Natl Acad Sci U S A 118 (2021).

33. S. Yeaman, et al., Convergent local adaptation to climate in distantly related conifers. Science (1979) 353, 1431–1433 (2016).

34. M. Bohutínská, C. L. Peichel, Divergence time shapes gene reuse during repeated adaptation. Trends Ecol Evol 39, 396–407 (2024).

35. K. E. Lotterhos, S. Yeaman, J. Degner, S. Aitken, K. A. Hodgins, Modularity of genes involved in local adaptation to climate despite physical linkage. Genome Biol 19, 1–24 (2018).

36. J. R. Whiting, et al., The genetic architecture of repeated local adaptation to climate in distantly related plants. Nat Ecol Evol (2024). 10.1038/s41559-024-02514-5.

37. Y. Duan, et al., MbICE1 Confers Drought and Cold Tolerance through Up-Regulating Antioxidant Capacity and Stress-Resistant Genes in Arabidopsis thaliana. Int J Mol Sci 23, 1–17 (2022).

38. J. S. Kim, S. Kidokoro, K. Yamaguchi-Shinozaki, K. Shinozaki, Regulatory networks in plant responses to drought and cold stress. Plant Physiol 195, 170–189 (2024).

39. R. Candido-Ribeiro, S. N. Aitken, Weak local adaptation to drought in seedlings of a widespread conifer. New Phytologist 241, 2395–2409 (2024).

40. J. Nuhu, “Genetic variation in drought and cold tolerance in selectively bred and natural populations of coastal Douglas-fir.” (2022).

41. S. Bansal, J. B. St. Clair, C. A. Harrington, P. J. Gould, Impact of climate change on cold hardiness of Douglas-fir (Pseudotsuga menziesii): Environmental and genetic considerations. Glob Chang Biol 21, 3814–3826 (2015).

42. G. E. Rehfeldt, Ecological adaptations in Douglas-Fir (Pseudotsuga menziesii var. glauca): a Synthesis. For Ecol Manage 28, 203–215 (1989).

43. P. F. Gugger, S. Sugita, J. Cavender-Bares, Phylogeography of Douglas-fir based on mitochondrial and chloroplast DNA sequences: Testing hypotheses from the fossil record. Mol Ecol 19, 1877–1897 (2010).

44. B. Lind, VarscanPipeline. (2021). Available at: https://github.com/CoAdapTree/varscan_pipeline/tree/v1.0.0 [Accessed 16 September 2022].

45. B. Lind, CMH-Test. (2023). Available at: https://github.com/brandonlind/cmh_test/tree/1.0.0 (URL) [Accessed 16 December 2023].

46. R. A. W. Wiberg, O. E. Gaggiotti, M. B. Morrissey, M. G. Ritchie, Identifying consistent allele frequency differences in studies of stratified populations. Methods Ecol Evol 8, 1899–1909 (2017).

47. N. Minadakis, et al., Polygenic architecture of flowering time and its relationship with local environments in the grass Brachypodium distachyon. Genetics 227, 1–17 (2024).

48. M. de Miguel, et al., Polygenic adaptation and negative selection across traits, years and environments in a long-lived plant species (Pinus pinaster Ait., Pinaceae). Mol Ecol 31, 2089–2105 (2022).

49. P. Singh, et al., Genetic architecture of disease resistance and tolerance in Douglas-fir trees. New Phytologist 243, 705–719 (2024).

50. I. R. MacLachlan, et al., Genome-wide shifts in climate-related variation underpin responses to selective breeding in a widespread conifer. Proc Natl Acad Sci U S A 118 (2021).

51. I. Höllinger, B. Wölfl, J. Hermisson, A theory of oligogenic adaptation of a quantitative trait. Genetics 225, 1–23 (2023).

52. G. Nocchi, J. R. Whiting, S. Yeaman, Repeated global adaptation across plant species. Proceedings of the National Academy of Sciences 121 (2024).

53. B. M. Lind, et al., How useful is genomic data for predicting maladaptation to future climate? Glob Chang Biol 30, 1–19 (2024).

54. K. J. Liepe, A. Hamann, P. Smets, C. R. Fitzpatrick, S. N. Aitken, Adaptation of lodgepole pine and interior spruce to climate: Implications for reforestation in a warming world. Evol Appl 9, 409–419 (2016).

55. V. Le Corre, A. Kremer, Genetic Variability at Neutral Markers, Quantitative Trait Loci and Trait in a Subdivided Population Under Selection. Genetics 164, 1205–1219 (2003).

56. I. R. MacLachlan, T. Wang, A. Hamann, P. Smets, S. N. Aitken, Selective breeding of lodgepole pine increases growth and maintains climatic adaptation. For Ecol Manage 391, 404–416 (2017).

57. M. Mimura, S. N. Aitken, Adaptive gradients and isolation-by-distance with postglacial migration in Picea sitchensis. Heredity (Edinb) 99, 224–232 (2007).

58. I. R. MacLachlan, S. Yeaman, S. N. Aitken, Growth gains from selective breeding in a spruce hybrid zone do not compromise local adaptation to climate. Evol Appl 11, 166–181 (2018).

59. D. Montwé, M. Isaac-Renton, A. Hamann, H. Spiecker, Cold adaptation recorded in tree rings highlights risks associated with climate change and assisted migration. Nat Commun 9, 1–7 (2018).

60. G. T. Howe, et al., From genotype to phenotype: Unraveling the complexities of cold adaptation in forest trees. Canadian Journal of Botany 81, 1247–1266 (2003).

61. S. Bansal, C. A. Harrington, P. J. Gould, J. B. St.Clair, Climate-related genetic variation in drought-resistance of Douglas-fir (Pseudotsuga menziesii). Glob Chang Biol 21, 947–958 (2015).

62. M. Isaac-Renton, et al., Northern forest tree populations are physiologically maladapted to drought. Nat Commun 9, 5254 (2018).

63. G. E. Rehfeldt, Genetic Differentiation of Douglas-Fir Populations from the Northern Rocky Mountains. Ecology 59, 1264–1270 (1978).

64. G. E. Rehfeldt, Development and Verification of Models of Freezing Tolerance for Douglas-fir Populations in the Inland Northwest. USDA For. Serv. Intermountain Res. Stn., Res. Pap. INT-369 2, 5 pp. (1986).

65. O. Savolainen, M. Lascoux, J. Merilä, Ecological genomics of local adaptation. Nat Rev Genet 14, 807–820 (2013).

66. O. Savolainen, T. Pyhäjärvi, T. Knürr, Gene Flow and Local Adaptation in Trees. Annu Rev Ecol Evol Syst 38, 595–619 (2007).

67. A. Kremer, et al., Long-distance gene flow and adaptation of forest trees to rapid climate change. Ecol Lett 15, 378–392 (2012).

68. E. L. Berdan, et al., Genomic architecture of supergenes: Connecting form and function. Philosophical Transactions of the Royal Society B: Biological Sciences 377 (2022).

69. L. H. Rieseberg, Chromosomal rearrangements and speciation. Trends Ecol Evol 16, 351–358 (2001).

70. M. Todesco, et al., Massive haplotypes underlie ecotypic differentiation in sunflowers. Nature 584, 602–607 (2020).

71. P. Battlay, et al., Large haploblocks underlie rapid adaptation in the invasive weed Ambrosia artemisiifolia. Nat Commun 14 (2023).

72. E. Moran, J. Lauder, C. Musser, A. Stathos, M. Shu, The genetics of drought tolerance in conifers. New Phytologist 216, 1034–1048 (2017).

73. C. Y. Y. Chang, K. Bräutigam, N. P. A. Hüner, I. Ensminger, Champions of winter survival: cold acclimation and molecular regulation of cold hardiness in evergreen conifers. New Phytologist 229, 675–691 (2021).

74. M. E. James, T. Brodribb, I. J. Wright, L. H. Rieseberg, D. Ortiz-Barrientos, Replicated Evolution in Plants. Annu Rev Plant Biol 74, 697–725 (2023).

75. N. Barghi, J. Hermisson, C. Schlötterer, Polygenic adaptation: a unifying framework to understand positive selection. Nat Rev Genet 21, 769–781 (2020).

76. N. McDowell, et al., Mechanisms of plant survival and mortality during drought: Why do some plants survive while others succumb to drought? New Phytologist 178, 719–739 (2008).

77. T. J. Brodribb, S. A. M. McAdam, G. J. Jordan, S. C. V. Martins, Conifer species adapt to low-rainfall climates by following one of two divergent pathways. Proc Natl Acad Sci U S A 111, 14489–14493 (2014).

78. R. Dauwe, J. A. Holliday, S. N. Aitken, S. D. Mansfield, Metabolic dynamics during autumn cold acclimation within and among populations of Sitka spruce (Picea sitchensis). New Phytologist 194, 192–205 (2012).

79. G. T. Howe, K. Jayawickrama, M. Cherry, G. R. Johnson, N. C. Wheeler, “Breeding Douglas-fir” in Plant Breeding Reviews. Vol. 27, 1st Ed., J. Janick, Ed. (WILEY, 2006).

80. S. Bansal, C. A. Harrington, J. B. St. Clair, Tolerance to multiple climate stressors: A case study of Douglas-fir drought and cold hardiness. Ecol Evol 6, 2074–2083 (2016).

81. A. Chaves-Sanjuan, et al., Structural basis of the regulatory mechanism of the plant CIPK family of protein kinases controlling ion homeostasis and abiotic stress. Proc Natl Acad Sci U S A 111, E4532–E4541 (2014).

82. L. Wang, et al., Arabidopsis UBC13 differentially regulates two programmed cell death pathways in responses to pathogen and low-temperature stress. New Phytologist 221, 919–934 (2019).

83. T. Tsuzuki, K. Takahashi, M. Tomiyama, S. I. Inoue, T. Kinoshita, Overexpression of the Mg-chelatase H subunit in guard cells confers drought tolerance via promotion of stomatal closure in Arabidopsis thaliana. Front Plant Sci 4, 1–8 (2013).

84. R. Nishiyama, et al., Analysis of cytokinin mutants and regulation of cytokinin metabolic genes reveals important regulatory roles of cytokinins in drought, salt and abscisic acid responses, and abscisic acid biosynthesis. Plant Cell 23, 2169–2183 (2011).

85. J. B. St Clair, N. L. Mandel, K. W. Vance-Borland, Genecology of Douglas fir in Western Oregon and Washington. Ann Bot 96, 1199–1214 (2005).

86. J. Paulose, O. Hallatschek, The impact of long-range dispersal on gene surfing. Proc Natl Acad Sci U S A 117, 7584–7593 (2020).

87. B. M. Lind, et al., How useful is genomic data for predicting maladaptation to future climate? Glob Chang Biol 30, 1–19 (2024).

88. J. Archambeau, et al., Evaluating genomic offset predictions in a forest tree with high population genetic structure. bioRxiv 2005–2024 (2024).

89. E. H. Murchie, T. Lawson, Chlorophyll fluorescence analysis: A guide to good practice and understanding some new applications. J Exp Bot 64, 3983–3998 (2013).

90. D. G. Butler, B. R. Cullis, A. R. Gilmour, B. J. Gogel, R. Thompson, ASReml-R Reference Manual Version 4. ASReml-R Reference Manual 176 (2018).

91. J. Pinheiro, D. Bates, S. Debroy, Sarkar D, R Core Team, nlme: Linear and Nonlinear Mixed Effects Models. R package version 3.1-117. J Apic Res 19, 196–199 (2014).

92. M. Hannerz, S. N. Aitken, J. N. King, S. Budge, Effects of genetic selection for growth on frost hardiness in western hemlock. Canadian Journal of Forest Research-Revue Canadienne De Recherche Forestiere 29, 509–516 (1999).

93. S. Ferrari, F. Cribari-Neto, Beta Regression for Modelling Rates and Proportions. J Appl Stat 31, 799–815 (2004).

94. F. Cribari-Neto, A. Zeileis, Beta Regression in R. J Stat Softw 34, 129–150 (2010).

95. J. C. Douma, J. T. Weedon, Analysing continuous proportions in ecology and evolution: A practical introduction to beta and Dirichlet regression. Methods Ecol Evol 10, 1412–1430 (2019).

96. C. Schlötterer, R. Tobler, R. Kofler, V. Nolte, Sequencing pools of individuals — mining genome-wide polymorphism data without big funding. Nat Rev Genet 15, 749–763 (2014).

97. B. M. Lind, et al., Haploid, diploid, and pooled exome capture recapitulate features of biology and paralogy in two non-model tree species. Mol Ecol Resour 22, 225–238 (2022).

98. R Core Team, A language and environment for statistical computing. (2021).

99. R. A. W. Wiberg, O. E. Gaggiotti, M. B. Morrissey, M. G. Ritchie, Identifying consistent allele frequency differences in studies of stratified populations. Methods Ecol Evol 8, 1899–1909 (2017).

100. D. B. Neale, et al., The Douglas-Fir genome sequence reveals specialization of the photosynthetic apparatus in Pinaceae. G3: Genes, Genomes, Genetics 7, 3157–3167 (2017).

101. A. Alexa, J. Rahnenführer, Gene set enrichment analysis with topGO. R package v.2.56.0 27 (2024).

102. M. Gautier, R. Vitalis, L. Flori, A. Estoup, f-Statistics estimation and admixture graph construction with Pool-Seq or allele count data using the R package poolfstat. Mol Ecol Resour 22, 1394–1416 (2022).

103. T. Wang, A. Hamann, D. Spittlehouse, C. Carroll, Locally downscaled and spatially customizable climate data for historical and future periods for North America. PLoS One 11, 1307–1309 (2016).

